# Phasing and imputation of single nucleotide polymorphism data of missing parents of bi-parental plant populations

**DOI:** 10.1101/2020.02.07.938795

**Authors:** Serap Gonen, Valentin Wimmer, R. Chris Gaynor, Ed Byrne, Gregor Gorjanc, John M. Hickey

## Abstract

This paper presents an extension to a heuristic method for phasing and imputation of genotypes of descendants in bi-parental populations so that it can phase and impute genotypes of parents of bi-parental populations that are fully ungenotyped or partially genotyped. The imputed genotypes of the parent are then used to impute low-density genotyped descendants of the bi-parental population to high-density. The extension works in three steps. First, it identifies whether a parent has no or low-density genotypes available and it identifies all of its relatives that have high-density genotypes. Second, using the high-density information of relatives, it determines whether the parent is homozygous or heterozygous for a given locus. Third, it phases heterozygous positions of the parent by matching haplotypes to its relatives.

We implemented the new algorithm in an extension of the AlphaPlantImptue software and tested its accuracy of imputing missing parent genotypes in simulated bi-parental populations from different scenarios. We also tested the accuracy of imputation of the missing parent’s descendants using the true genotype of the parent and compared this to using the imputed genotypes of the parent. Our results show that across all scenarios, the accuracy of imputation of a parent, measured as the correlation between true and imputed genotypes, was > 0.98 and did not drop below ∼ 0.96. The imputation accuracy of a parent was always higher when it was inbred than when it was outbred and when it had low-density genotypes. Including ancestors of the parent at HD, increasing the number of crosses and the number of high-density descendants all increased the accuracy of imputation. The high imputation accuracy achieved for the parent across all scenarios translated to little or no impact on the accuracy of imputation of its descendants at low-density.

**Key Message:** New fast and accurate method for phasing and imputation of SNP chip genotypes within diploid bi-parental plant populations.

## Introduction

This paper presents an extension to a heuristic method for phasing and imputation of genotypes of descendants in bi-parental populations so that it can phase and impute genotypes of parents of bi-parental populations that are fully ungenotyped or partially genotyped. The imputed genotypes of the parent are then used to impute low-density genotyped descendants of the bi-parental population to high-density. High-density SNP array data in plant breeding populations is increasingly valuable for genomic selection and for identifying regions of the genome that underlie traits of interest in genome-wide association studies (Bernardo and Yu, 2007; Hamblin et al., 2011). One of the major barriers to the adoption of genomic selection in plant breeding programs is that the number of selection candidates that would need to be genotyped at high-density in each cycle can be very large (Heffner et al., 2010).

In livestock and human populations, an effective strategy to overcome this cost barrier has been to genotype a subset of the population at high-density and to use this data for imputation of the rest of the population genotyped at low-density. The adoption of this strategy has been enabled by the development of imputation tools that leverage pedigree relationships or population-level linkage information for fast and accurate genotype imputation (Kong et al., 2008; Howie et al., 2009; Druet and Georges, 2010; Li et al., 2010; Sargolzaei et al., 2011; Hickey et al., 2011; Cleveland and Hickey, 2013; Hickey and Kranis, 2013; VanRaden et al., 2015; O’Connell et al., 2016; Loh et al., 2016; Antolín et al., 2017).

In most plant breeding populations, a small number of selected parents are crossed to generate large numbers of bi-parental populations. Therefore, high-density genotyping of all parents and low-density genotyping of focal individuals (i.e., descendants that are the imputation targets) could be an effective low-cost strategy in these populations (Jacobson et al., 2014, 2015; Gorjanc et al., 2017b; a). To our knowledge, very few imputation tools designed to leverage features of plant breeding programs, such as fully or almost fully inbred parents, small numbers of meiosis separating parents and descendants who are to have genotypes imputed and different crossing structures (e.g., selfing, double haploids), to enable fast and accurate genotype imputation have been developed. We recently presented a fast, computationally efficient and accurate heuristic genotype imputation method implemented in AlphaPlantImpute (Gonen et al., 2018) that explicitly leverages features of plant breeding programs to maximise the accuracy of imputation. Using simulated data, we showed that an average accuracy of imputation of 0.96 could be achieved for a scenario where F_2_ individuals who were to be imputed were genotyped with 50 markers per chromosome and both parents were inbred and genotyped at 25,000 markers per chromosome.

The drawback of our previous algorithm is that it requires that both parents of each bi-parental population are known and have phased genotypes available at high-density. Although this is normally the case when parents are inbred, pedigree errors, sample loss or mislabelling or poor DNA quality can mean that one or both parents may have fully or partially missing genotype data. Additionally, if genotyping resources are limiting, breeders may choose not to genotype a parent that has only been used to in one or two crosses. Furthermore, even if parents have high-density genotypes available, unless they are fully inbred (i.e., homozygous at every locus and therefore all genotypes are phased *de facto*) it is unlikely that they have phased genotypes available for use in imputation.

This paper presents an extension to our previous algorithm in AlphaPlantImpute to enable it phase and impute high-density genotypes of parents of bi-parental populations that are missing or that only have low-density genotypes available. The extension requires that some relatives of the parent (e.g., descendants, ancestors, siblings) have high-density genotypes. The extension has three steps. First, it identifies whether a parent has no or low-density genotypes available and all of its relatives that have high-density genotypes. Second, using the high-density information of relatives, it determines whether the parent is homozygous or heterozygous for a given locus. Third, it phases heterozygous positions of the parent by matching haplotypes to its relatives.

We tested the accuracy of imputing missing parent genotypes using the extension to AlphaPlantImpute in simulated bi-parental populations from different scenarios. These scenarios varied in the levels of inbreeding in the missing parent, whether the parent had no genotypes or was genotyped at low-density, the number of crosses that the parent was used in and whether the ancestors of the parent had high-density genotypes available. We calculated the accuracy of imputation of the missing parent within each scenario as the correlation between the true and imputed genotypes. We also tested the accuracy of imputation of the missing parent’s descendants using the true genotype of the parent compared to using the imputed genotypes of the parent. Our results show that across all scenarios, the accuracy of imputation of a parent was consistently high. The imputation accuracy of a parent was always higher when it was inbred than when it was outbred and when it had low-density genotypes. Including ancestors of the parent at HD, increasing the number of crosses and increasing the number of high-density descendants all increased the accuracy of imputation. The high imputation accuracy achieved for the parent across all scenarios had little or no impact on the accuracy of imputation of its descendants at low-density, which remained high.

## Materials and methods

### Definitions

A focal individual is a descendant individual that is to be imputed. Parent A is the missing parent that is the target of imputation. The high-density (**HD**) array is the target array for imputation. In our test datasets, the HD array consisted of 25,000 SNP markers. The low-density (**LD**) array is the array at which focal individuals have genotypes and where Parent A may have genotypes. The LD array consisted of 50 SNP markers.

### Description of the method

We present an extension to the original imputation method in AlphaPlantImpute to phase and impute parents of bi-parental populations that are missing or that have LD genotypes available. First, AlphaPlantImpute identifies parents with missing genotypes or unphased genotypes (hereafter described for a single parent referred to as Parent A). Second, AlphaPlantImpute gathers HD genotype information of all known relatives for Parent A. Relatives include ancestors, siblings, descendants and mates. AlphaPlantImpute then uses any genotype information available on Parent A and its relatives to first impute missing genotypes and then phase heterozygous genotypes of Parent A.

### Parent A not genotyped

In livestock, the next generation are produced by a single cross of two ancestors. This means that loci where both ancestors are homozygous for the same genotype (i.e., both are genotype 0 or genotype 2) and where ancestors are opposing homozygotes (i.e., one is genotype 0 and the other is 2) can be confidently imputed in their offspring. In plant breeding populations, individuals are often the product of a single cross to produce F1 individuals, followed by many rounds of selfing. This means that if an offspring (in this case Parent A) has no genotypes but has ancestors genotyped at HD, the only loci that can be confidently imputed are where both of its ancestors are homozygous for the same. These loci are phased *de-facto*.

If Parent A has HD descendants and mates, use this information to phase and impute genotypes for Parent A in the following three steps: (1) Infer positions where Parent A is likely to be homozygous based on allele frequencies in descendants. For example, if all HD descendants are fixed for the 0 allele, then Parent A is likely to be genotype 0. If the allele frequencies are almost equal and the mate of Parent A is known to be genotype 0, then Parent A is likely to be genotype 2; (2) Infer positions where Parent A is likely to be heterozygous based on genotype frequency distortion in descendants. This is calculated using a chi-square test of observed genotype counts to expected genotype counts given observed allele frequencies. If there is significant distortion and the mate is homozygous then Parent A is likely to be heterozygous; (3) To phase inferred heterozygous loci of Parent A at HD, collate the genotypes of all HD descendants and mates at these loci. Use these loci as anchor points in the heuristic imputation algorithm of AlphaPlantImpute (Gonen et al., 2018) to determine parent-of-origin for the haplotypes of all descendants. For haplotypes of descendants assigned to Parent A, collate the haplotypes at HD and derive consensus phase for Parent A.

### Parent A has LD genotypes

If Parent A has LD genotypes and has ancestors genotyped at HD, AlphaPlantImpute uses the LD genotypes in the heuristic imputation algorithm as described in Gonen et. al. (2018). Briefly, the LD genotypes serve as anchor points for defining parent-of-origin for the haplotypes of Parent A. Use these anchor points to simultaneously phase and impute Parent A to HD.

If Parent A has HD descendants and mates, impute the genotypes of Parent A in the following four steps: (1) Identify the loci at which Parent A, descendants and mates are genotyped and collate the genotypes; (2) Use these genotypes as anchor points in the existing heuristic imputation algorithm of AlphaPlantImpute (Gonen et al., 2018) to determine parent-of-origin for the haplotypes of all descendants; (3) For haplotypes of descendants assigned to Parent A, collate the haplotypes at HD and derive consensus haplotypes for Parent A; (4) Fill genotypes of Parent A as the sum of the two derived haplotypes.

If Parent A has HD ancestors, descendants and mates then a consensus of the phased and imputed genotypes using only ancestor information or using only descendant information is derived. Where they disagree, set as missing.

### Examples of implementation: Description of datasets

To test the imputation accuracy of this modification of AlphaPlantImpute, testing datasets of bi-parental populations from different scenarios were simulated. These scenarios varied in the levels of inbreeding in the missing parent, whether the parent had no genotypes or was genotyped at low-density, the number of crosses that the parent was used in and whether the ancestors of the parent had high-density genotypes available. A description of the general structure and simulation method of the different scenarios is given below.

### Simulation of genomic data

Sequence data for 100 base haplotypes for a single chromosome were simulated using the Markovian Coalescent Simulator (Chen et al., 2009) and AlphaSimR (Faux et al., 2016). The base haplotypes were 10^8^ base pairs in length, with a per site mutation rate of 1.0×10^−8^ and a per site recombination rate of 1.0×10^−8^, resulting in a chromosome size of 1 Morgan (M). The effective population size (N_e_) was set at specific points during the simulation to mimic changes in N_e_ in a crop such as maize *(Zea mays L*.). These set points were: 100 in the base generation, 1000 at 100 generations ago, and 10,000 at 2000 generations ago, with linear changes in between. The resulting whole-chromosome haplotypes had approximately 80,000 segregating sites in total.

### Simulation of a pedigree

A founder population of 1000 inbred individuals was initiated. Two individuals from this founder population (denoted B and C) were crossed to generate 1000 F_1_ individuals. These individuals were selfed for *n* rounds and one individual was selected to be Parent A. The number of rounds of selfing (*n*) was 100 if Parent A was simulated to be fully inbred or was 1 if Parent A was simulated to be outbred. Depending on the scenario, Parent A was crossed to 1, 2, 3 or 4 individuals (denoted D, E, F, G) from the initial founder population to generate 1000 of F_1_ individuals. F_1_ individuals were selfed to generate 1000 F_2_ individuals. These were the descendants used for imputation of Parent A.

In the base generation, individuals had their chromosomes sampled from the 100 base haplotypes. In subsequent generations the chromosomes of each individual was sampled from parental chromosomes with recombination, resulting in a chromosome size of 1 Morgan (M). Recombinations occurred with a 1% probability per cM and were uniformly distributed along the chromosome.

### Simulated SNP marker arrays

A single HD array of 5,000 SNP markers and a single LD array of 50 SNP markers for the single chromosome was simulated. Arrays were constructed by aiming to select a set of markers that segregated in the parents and that were evenly distributed across the chromosome. The LD array was nested within the HD array.

### Scenarios

The imputation accuracy of Parent A was assessed in 8 different scenarios. Scenarios were designed to test the effect of including or excluding ancestors of Parent A (hereafter referred to as Grandparent 1 and Grandparent 2) and the effect of having genotype information of F_2_ individuals from one, two, three or four crosses of Parent A with Parents B, C, D and E. From each cross, 10 F_2_ individuals were selected as HD descendants. The remaining 990 were F_2_ focal individuals genotyped at LD. In all scenarios, Parent A could be either inbred or outbred and could be either genotyped at LD or not. One hundred replications of each scenario were performed and the average of each replication is reported in the results.

Scenarios 1, 2, 3 and 4 excluded the parents of Parent A (hereafter referred to as Grandparent 1 and Grandparent 2). Scenarios 5, 6, 7 and 8 included Grandparent 1 and Grandparent 2. Scenarios 1 and 5 had information from one cross (Parent A x Parent B). Scenarios 2 and 6 had information from two crosses (Parent A x Parent B; Parent A x Parent C). Scenarios 3 and 7 had information from three crosses (Parent A x Parent B; Parent A x Parent C; Parent A x Parent D). Scenarios 4 and 8 had information from three crosses (Parent A x Parent B; Parent A x Parent C; Parent A x Parent D; Parent A x Parent E).

In addition to the imputation accuracy of Parent A, the accuracy of imputing the F_2_ focal individuals genotyped at LD to HD using the phased and imputed genotypes of Parent A was assessed. This was compared to the imputation accuracy that would have been achieved if genotypes of Parent A were known and not imputed.

### Analysis

Imputation of Parent A was performed using information across all crosses and of Parents B and C, if available. Imputation of F_2_ focal individuals genotyped at LD was performed within a cross using the heuristic imputation method of AlphaPlantImpute described in Gonen et. al. 2018. The imputation accuracy was calculated as the correlation between the true and imputed genotypes. The imputation yield was calculated as the number of SNPs with imputed genotypes divided by the total number of SNPs on the HD array. In all scenarios, Grandparents 1 and 2 and Parents B, C, D and E were assumed genotyped at HD.

## Results

Unless otherwise stated, all results presented below had 10 HD descendants per cross.

### Effect of whether Parent A is inbred or outbred

The imputation accuracy of Parent A was always higher when it was inbred than when it was outbred but the differences were small. Figure 1 plots the genotype accuracy for Parent A in Scenario 1. The colours differentiate whether Parent A was inbred (red) or outbred (blue). The transparencies differentiate whether Parent A had no genotypes (opaque) or had LD genotypes (transparent). Figure 1 shows that when Parent A had no genotypes, the accuracy of imputation was 1.01 times higher when it was inbred than when it was outbred (0.980 vs. 0.970). When Parent A had LD genotypes, the accuracy of imputation was 1.02 times higher when it was inbred than when it was outbred (0.999 vs. 0.983). For all cases, the yield of imputation was 100%.

**Figure 1.**
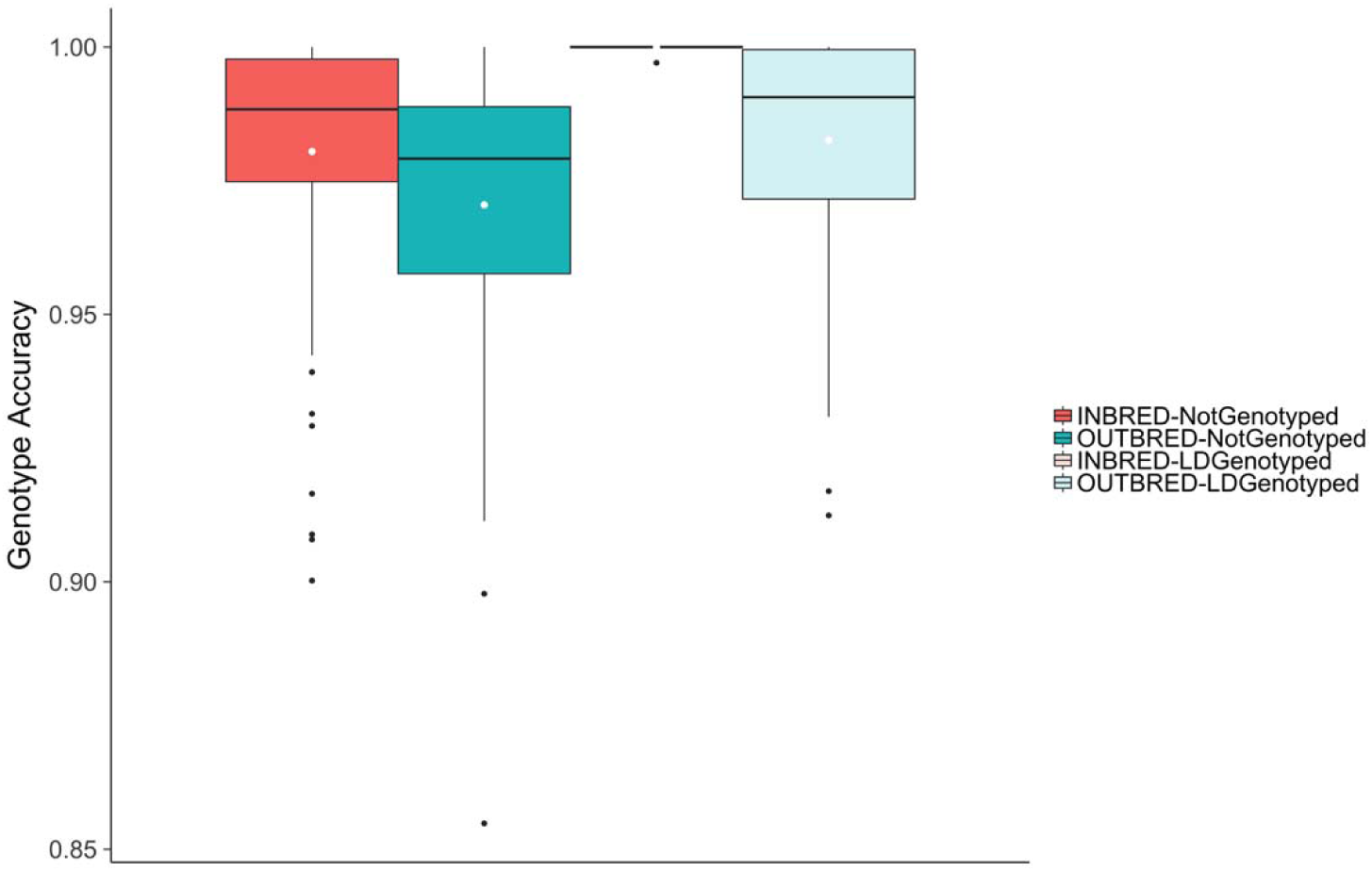
Effect of whether Parent A is inbred or outbred and whether Parent A has no or LD genotypes. Genotype imputation accuracy for Parent A in Scenario 1. The colours differentiate whether Parent A was inbred (red) or outbred (blue). The transparencies differentiate whether Parent A had no genotypes (opaque) or had LD genotypes (transparent).

### Effect of whether Parent A has LD genotypes or not

The imputation accuracy of Parent A was always higher when it had LD genotypes than when it had no genotypes but the differences were small. Figure 1 shows that when Parent A was inbred, the accuracy of imputation was 1.02 times higher when it had LD genotypes than when it had no genotypes (0. 999 vs. 0. 980). When Parent A was outbred, the accuracy of imputation was 1.01 times higher when it had LD genotypes than when it had no genotypes but the differences were small (0. 983 vs. 0.970).

### Effect of including Grandparent 1 and Grandparent 2 at HD

Including Grandparent 1 and Grandparent 2 increased the accuracy of imputation when Parent A has some LD genotypes but the differences were small. When Parent A had no genotypes, the accuracy of imputation was the same regardless of whether Grandparent 1 and Grandparent 2 were included or excluded. Figure 2 is similar to Figure 1 and plots the genotype accuracy (Figure 2a) and genotype yield (Figure 2b) for Parent A in Scenarios 1 and 5. Figure 2a shows that the main benefit of including Grandparent 1 and Grandparent 2 for increasing the imputation accuracy was when Parent A was outbred and had LD genotypes. In this case, the accuracy of imputation of Parent A was 1.02 times higher when Grandparent 1 and Grandparent 2 were included than when they were excluded (0.983 vs. 0.997). However, this increase in accuracy was at the expense of yield. Figure 2b shows that when Parent A was outbred and had LD genotypes, the yield was 100% when Grandparent 1 and Grandparent 2 were excluded and was 97.4% when Grandparent 1 and Grandparent 2 were included.

**Figure 2.**
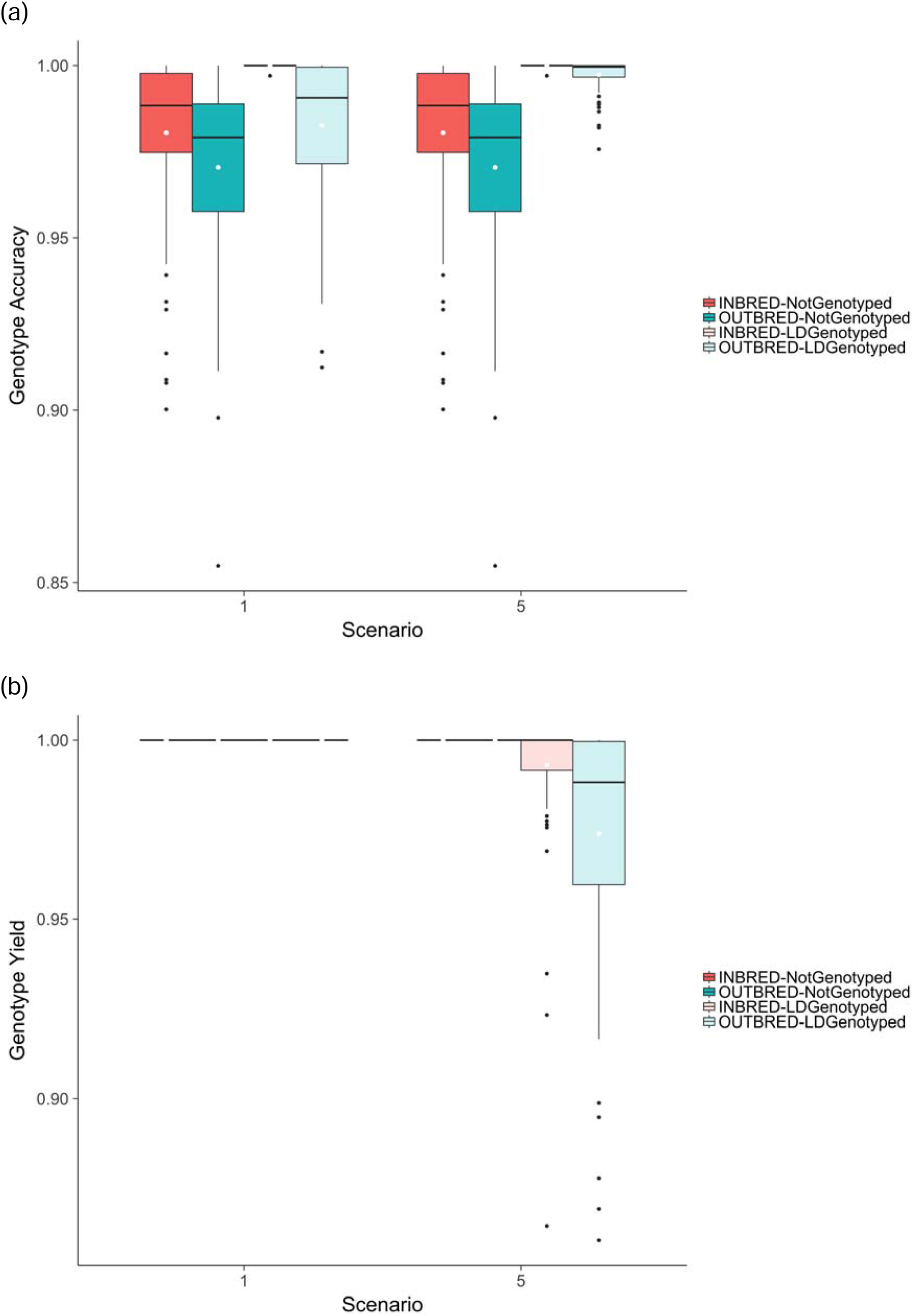
Effect of including ancestors of Parent A at HD. Genotype imputation accuracy (a) and imputation yield (b) for Parent A in Scenarios 1 and 5. The colours differentiate whether Parent A was inbred (red) or outbred (blue). The transparencies differentiate whether Parent A had no genotypes (opaque) or had LD genotypes (transparent).

### Effect of the number of crosses with Parent A

Increasing the number of crosses that Parent A was used in increased the accuracy of imputation but the differences were small. Figure 3a is similar to Figure 1 and plots the genotype accuracy for Parent A in Scenarios 1, 2, 3 and 4. Figure 3a shows that increasing the number of crosses from one in Scenario 1 to two in Scenario 2 increased the imputation accuracy regardless of whether Parent A was inbred or outbred, or had no genotypes or had LD genotypes. When Parent A was inbred, the accuracy of imputation was 1.02 times higher in Scenario 2 than in Scenario 1 when it had no genotypes (0.980 vs. 0.999) and was just slightly higher when it had LD genotypes (0.999 vs. 1.0). When Parent A was outbred, the accuracy of imputation was 1.01 times higher in Scenario 2 than in Scenario 1 when it had no genotypes (0.970 vs. 0.975) and was 1.01 times higher when it had LD genotypes (0.983 vs. 0.992). For all cases, the yield of imputation was 100%.

**Figure 3.**
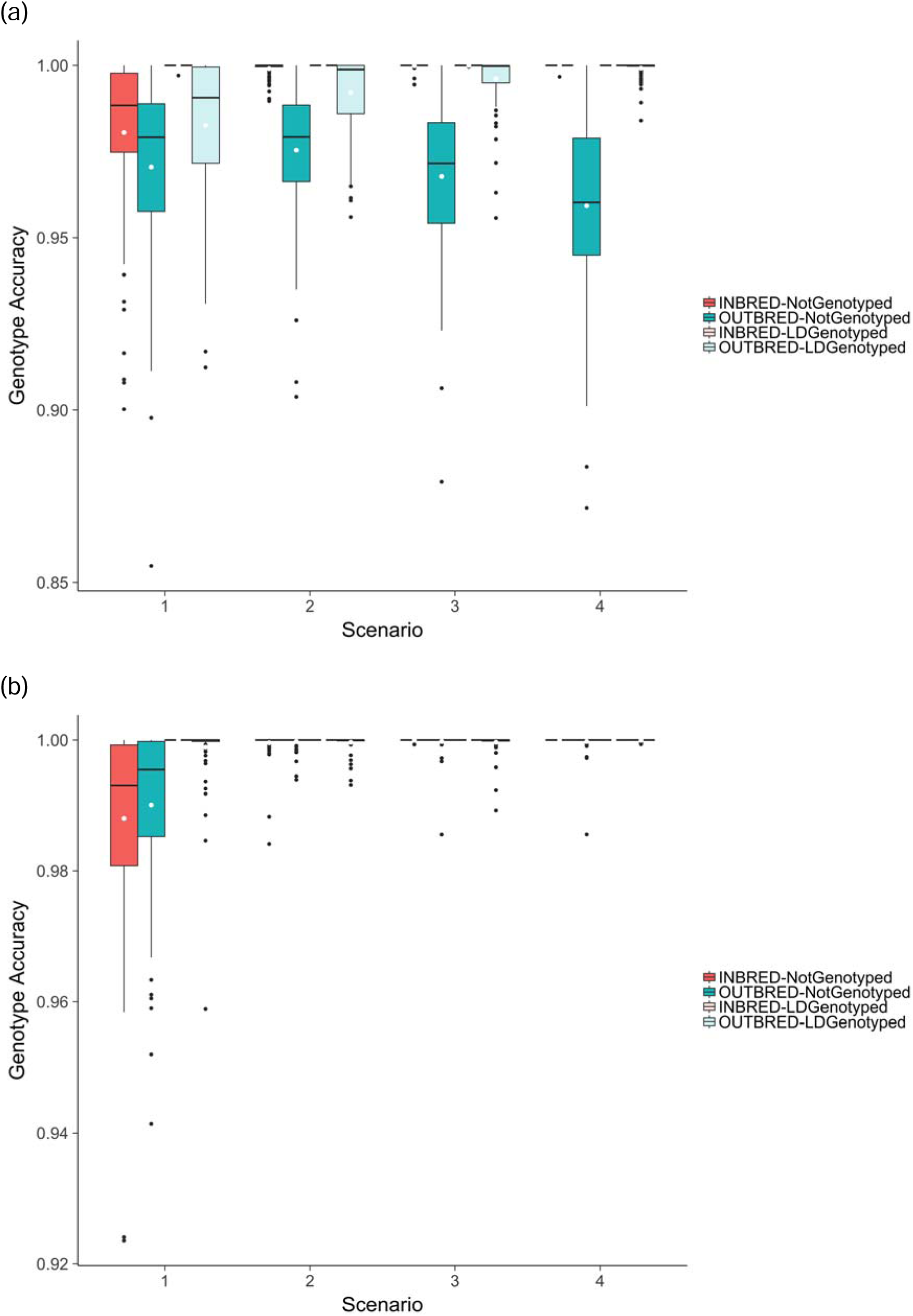
Effect of the number of crosses and number of HD descendants per cross. Genotype imputation accuracy for Parent A with 10 HD descendants per cross (a) and with 50 HD descendants per cross (b) in Scenarios 1, 2, 3 and 4. The colours differentiate whether Parent A was inbred (red) or outbred (blue). The transparencies differentiate whether Parent A had no genotypes (opaque) or had LD genotypes (transparent).

Increasing the number of crosses that Parent A was used in increased the accuracy of imputation most when Parent A was outbred and had LD genotypes but the differences were small. Figure 3a shows that when the number of crosses increased from one in Scenario 1 to four in Scenario 4, the accuracy of imputation was 1.02 times higher in Scenario 4 than in Scenario 1 when Parent A was outbred and had LD genotypes (0.983 vs. 0.999).

Figure 3a also shows that increasing the number of crosses that Parent A was used in decreased the accuracy of imputation when Parent A was outbred and had no genotypes but the differences were small. When the number of crosses increased from one in Scenario 1 to four in Scenario 4, the accuracy of imputation was 1.01 times higher in Scenario 1 than in Scenario 4 (0.970 vs. 0.959).

### Effect of number of descendants with HD genotypes

Increasing the number of descendants with HD genotypes increased the accuracy of imputation of Parent A but the differences were small. Figure 3b is similar to Figure 3a and plots the genotype accuracy for Parent A in Scenarios 1, 2, 3 and 4 when the number of descendants with HD genotypes was 50. For example for Scenario 1, when the number of descendants increased from 10 to 50 the accuracy of imputation was 1.01 times higher when Parent A was inbred and had no genotypes (0.980 vs. 0.988), was just slightly higher when Parent A was inbred and had LD genotypes (0.999 vs. 1.00), was 1.02 times higher when Parent A was outbred and had no genotypes (0.970 vs. 0.990), and was 1.02 times higher when Parent A was outbred and had LD genotypes (0.983 vs. 0.999). For all cases, the yield of imputation was 100%. Figure 3b also shows that when the number of descendants with HD genotypes was 50, increasing the number of crosses to two or more resulted in accuracy of imputation for Parent A of >0.999.

### Effect of using imputed genotypes or true genotypes of Parent A to impute F_2_ focal individuals

Using true or imputed genotypes of Parent A had only a small effect on the accuracy of imputation of impute F_2_ focal individuals. Figure 4 plots the increase in imputation accuracy achieved for F_2_ focal individuals for Scenario 1. The increase in imputation accuracy is the difference between the accuracy achieved using true or imputed genotypes for Parent A to impute focal individuals. Figure 4 shows that the increase in imputation accuracy achieved for focal individuals using true genotypes of Parent A compared to using imputed genotypes was minimal regardless of whether Parent A was inbred or outbred or had LD or no genotypes. The largest increase achieved was when Parent A was outbred and had no genotypes, where an increase of 0.029 was achieved. When Parent A was inbred and had LD genotypes, there was no increase in the accuracy of imputation of focal individuals when using true or imputed genotypes for Parent A.

**Figure 4.**
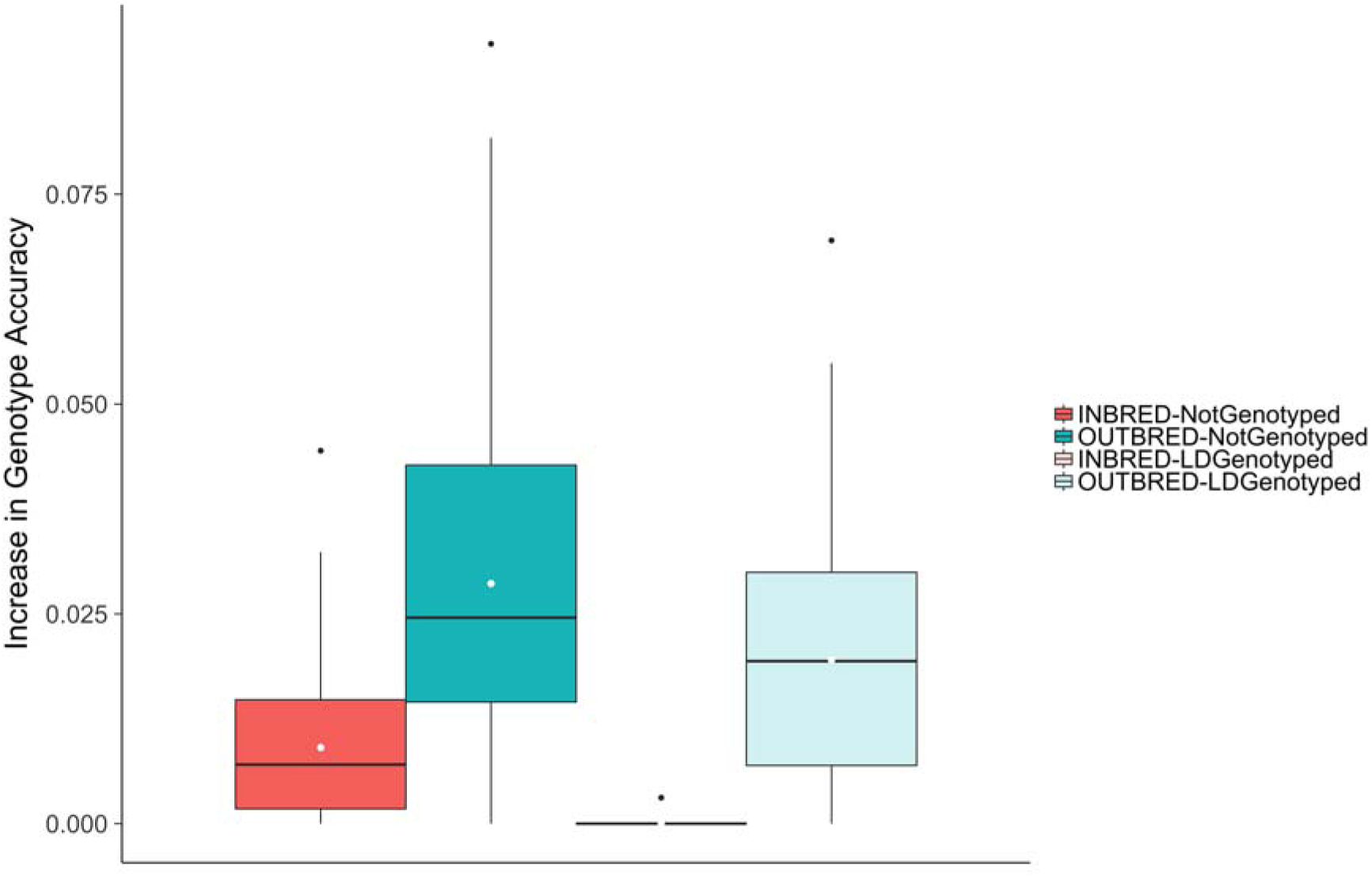
Effect of using imputed genotypes or true genotypes of Parent A to impute F_2_ focal individuals. Increase in the genotype imputation accuracy for F_2_ focal individuals using true rather than imputed genotypes for Parent A in Scenario 1. The colours differentiate whether Parent A was inbred (red) or outbred (blue). The transparencies differentiate whether Parent A had no genotypes (opaque) or had LD genotypes (transparent).

## Discussion

Our results highlight two main points for discussion: (i) the performance of AlphaPlantImpute in imputing Parent A; and (ii) the effect using imputed genotypes or true genotypes of Parent A to impute F_2_ focal individuals.

### Performance of AlphaPlantImpute in Imputing Parent A

This paper presents an extension to the original heuristic imputation method in AlphaPlantImpute (Gonen et al., 2018) to phase and impute genotypes for parents of bi-parental populations who are missing or who have LD genotypes available. The extension requires that some relatives of the parent (e.g., descendants, ancestors, siblings) have HD genotypes. We tested and compared the performance of the algorithm, which we implemented in an updated version of AlphaPlantImpute (Gonen et al., 2018), across a range of scenarios where the parent to be imputed (Parent A) could be inbred or outbred, could have no or LD genotypes, could be a parent of one or multiple crosses with descendants at HD, or could have parents with HD genotypes. In general across all scenarios, the average accuracy was > 0.98 and the average accuracy did not drop below ∼ 0.96. The yield was 100% for all scenarios apart from when Grandparents 1 and 2 (i.e., the ancestors of Parent A) were included with HD genotypes. The only scenario where this was not the case was when Grandparents 1 and 2 were included and Parent A was outbred and had LD genotypes. In this case, the yield dropped to 97%. The reason for this is that this scenario had HD genotypes available for both Grandparents 1 and 2 and for 10 offspring of Parent A. The heuristic algorithm uses the two sources of information independently to impute Parent A. Where they disagree, the genotype is set as missing.

As expected, adding more information from relatives genotyped at HD increased the accuracy of imputation for Parent A. When Parent A was used in a single cross, including its parents at HD increased the accuracy of imputation for Parent A, particularly when Parent A was outbred and had LD genotypes. However, the increase in accuracy when Parent A had LD genotypes was at the expense of yield. The reason for this decrease in yield is likely caused by disagreement between Parent A genotypes imputed using its descendants genotyped at HD and genotypes imputed using its parents genotyped at HD. When Parent A had no genotypes, including its parents at HD had no effect. This is because the only loci that could be filled with confidence were loci where its parents were fixed for the same allele.

Increasing the number of crosses that Parent A was used in increased the accuracy of imputation for Parent A when it was inbred or outbred and had LD genotypes. This was likely due to two reasons. First, the extra HD information from other crosses increased the ability to call heterozygous loci. For example, by chance within a single cross one of the haplotypes of Parent A may have been underrepresented or not represented in the descendants selected for HD genotyping but may have been represented in HD descendants in the second cross. Second, the LD genotypes of Parent A were used to assign parent-of-origin to the haplotypes of HD descendants. Loci that were not informative of parent-of-origin within one cross may have been informative in another cross, providing extra information on the haplotypes of Parent A. Increasing the number of crosses that Parent A was used in had only a small benefit when Parent A was inbred and had no genotypes. In this case, the accuracy of imputation for Parent A was already ∼ 0.98 with a single cross and increasing to number of crosses increased the accuracy of imputation for Parent A to > 0.999. The only exception to the benefit of increasing the number of crosses was when Parent A was outbred and had LD genotypes. This could have been caused by incorrect assignment or the inability to assign parent-of-origin to the haplotypes of HD descendants, which would result in incorrect or uncalled genotypes for Parent A.

Increasing the number of descendants at HD within a cross increased the accuracy of imputation across all scenarios. This is expected, since more HD relatives provides more information for confidently calling the genotypes of Parent A.

Overall, the results suggest that high imputation accuracy of >0.98 and an imputation yield of 100% in almost all cases can be achieved for Parent A by collating HD genotypes of as many relatives as possible. This is critical for ensuring accurate imputation of descendants genotyped at LD.

### Effect of using imputed genotypes or true genotypes of Parent A to impute F_2_ focal individuals

Using true or imputed genotypes of Parent A had only a small effect on the accuracy of imputation of impute F_2_ focal individuals. The largest increase in imputation accuracy when using true genotypes rather than imputed genotypes for Parent A was observed when Parent A was outbred and not genotyped, but even in this case the increase was 0.028. The likely reason for the small increase was that the accuracy of imputation of Parent A was in general > 0.96 across all scenarios. Therefore, our results suggest that some error in the imputation of Parent A is likely to have minimal, if any effect on the imputation of focal individuals that are its descendants.

### Relevance for breeding programs

The use of genomic information in plant breeding populations could have a large impact for informing selection decisions (Bernardo and Yu, 2007; Heffner et al., 2010; Hamblin et al., 2011; Hickey et al., 2014; Daetwyler et al., 2014; Bassi et al., 2016). However, the large cost associated with the large number of candidates that would need to be genotyped in order to leverage the power of genomic selection is still a bottleneck. One way of overcoming this bottleneck would be to genotype the many thousands of selection candidates at LD and impute them to HD. To do this, the parents of the candidates need to have phased HD genotypes available or inferred. Genotyping parents at HD and inferring phase is theoretically feasible. However, in practice, not all parents will have phased HD genotypes available due to: (1) low quality DNA samples; (2) missing DNA samples (for example for older samples); (3) parents that are used in only a single cross may not be worth genotyping; (4) incomplete pedigrees; and (5) pedigree errors. If relatives (e.g., ancestors, offspring, siblings or mates) of a parent have HD genotypes available, this information could be used to phase and impute HD genotypes for the missing parent. The imputed genotypes could then be used to impute any selection candidates that descend from this missing parent. Our simulations show that high imputation accuracy and yield can be obtained for a missing parent, providing a cost-effective and powerful way of obtaining accurate HD genotypes for selection candidates that are descendants of the imputed parent.

### Software availability

We implemented our method in a software package called AlphaPlantImpute, which is available for download at http://www.AlphaGenes.roslin.ed.ac.uk/AlphaPlantImpute/ along with a user manual.

## Conclusions

This paper presents an extension to a heuristic method implemented in AlphaPlantImptue so that it can phase and impute genotypes of parents of bi-parental populations that are fully ungenotyped or partially genotyped. The imputed genotypes of the parent are then used to impute low-density genotyped descendants of the bi-parental population to HD. Our results show that the imputation yield was 100% in almost all scenarios. The accuracy of imputation of a parent was > 0.98 and did not drop below ∼ 0.96. The imputation accuracy of a parent was always higher when it was inbred than when it was outbred and when it had low-density genotypes. Including ancestors of the parent at HD, increasing the number of crosses and increasing the number of high-density descendants all increased the accuracy of imputation. The high imputation accuracy achieved translated to little or no impact on the accuracy of imputation of its descendants at low-density, which remained high. This extension will be useful in plant breeding populations aiming to incorporate genomic selection for a large number of candidates genotyped at LD where one of the parents of those candidates has no HD phased genotypes available.

## Acknowledgments

The authors acknowledge financial support from the BBSRC ISP grant number ‘BB/P013759/1’ and from the BBSRC KWS grant number ‘BB/R002061/1’. This work has made use of the resources provided by the Edinburgh Compute and Data Facility (ECDF) (http://www.ecdf.ed.ac.uk).

## References

Antolín, R., C. Nettelblad, G. Gorjanc, D. Money, and J.M. Hickey. 2017. A hybrid method for the imputation of genomic data in livestock populations. Genet. Sel. Evol. 49(1): 30. doi: 10.1186/s12711-017-0300-y.

Bassi, F.M., A.R. Bentley, G. Charmet, R. Ortiz, and J. Crossa. 2016. Breeding schemes for the implementation of genomic selection in wheat (Triticum spp.). Plant Sci. 242: 23–36. doi: 10.1016/j.plantsci.2015.08.021.

Bernardo, R., and J. Yu. 2007. Prospects for Genomewide Selection for Quantitative Traits in Maize. Crop Sci. 47(3): 1082. doi: 10.2135/cropsci2006.11.0690.

Chen, G.K., P. Marjoram, and J.D. Wall. 2009. Fast and flexible simulation of DNA sequence data. Genome Res. 19(1): 136–142. doi: 10.1101/gr.083634.108.

Cleveland, M.A., and J.M. Hickey. 2013. Practical implementation of cost-effective genomic selection in commercial pig breeding using imputation. J. Anim. Sci. 91(8): 3583–3592. doi: 10.2527/jas.2013-6270.

Daetwyler, H.D., U.K. Bansal, H.S. Bariana, M.J. Hayden, and B.J. Hayes. 2014. Genomic prediction for rust resistance in diverse wheat landraces. Theor. Appl. Genet. 127(8): 1795–1803. doi: 10.1007/s00122-014-2341-8.

Druet, T., and M. Georges. 2010. A Hidden Markov Model Combining Linkage and Linkage Disequilibrium Information for Haplotype Reconstruction and Quantitative Trait Locus Fine Mapping. Genetics 184(3): 789–798. doi: 10.1534/genetics.109.108431.

Faux, A.-M., G. Gorjanc, R.C. Gaynor, M. Battagin, S.M. Edwards, et al. 2016. AlphaSim: Software for Breeding Program Simulation. Plant Genome 9(3). doi: 10.3835/plantgenome2016.02.0013.

Gonen, S., V. Wimmer, R.C. Gaynor, E. Byrne, G. Gorjanc, et al. 2018. A heuristic method for fast and accurate phasing and imputation of single-nucleotide polymorphism data in bi-parental plant populations. Theor. Appl. Genet. 131(11): 2345–2357. doi: 10.1007/s00122-018-3156-9.

Gorjanc, G., M. Battagin, J.-F. Dumasy, R. Antolin, R.C. Gaynor, et al. 2017a. Prospects for Cost-Effective Genomic Selection via Accurate Within-Family Imputation. Crop Sci. 57(1): 216. doi: 10.2135/cropsci2016.06.0526.

Gorjanc, G., J.-F. Dumasy, S. Gonen, R.C. Gaynor, R. Antolin, et al. 2017b. Potential of Low-Coverage Genotyping-by-Sequencing and Imputation for Cost-Effective Genomic Selection in Biparental Segregating Populations. Crop Sci. 57(3): 1404–1420. doi: 10.2135/cropsci2016.08.0675.

Hamblin, M.T., E.S. Buckler, and J.-L. Jannink. 2011. Population genetics of genomics-based crop improvement methods. Trends Genet. TIG 27(3): 98–106. doi: 10.1016/j.tig.2010.12.003.

Heffner, E.L., A.J. Lorenz, J.-L. Jannink, and M.E. Sorrells. 2010. Plant Breeding with Genomic Selection: Gain per Unit Time and Cost. Crop Sci. 50(5): 1681. doi: 10.2135/cropsci2009.11.0662.

Hickey, J.M., S. Dreisigacker, J. Crossa, S. Hearne, R. Babu, et al. 2014. Evaluation of genomic selection training population designs and genotyping strategies in plant breeding programs using simulation. Crop Sci. 54: 1476–1488. doi: 10.2135/cropsci2013.03.0195.

Hickey, J.M., B.P. Kinghorn, B. Tier, J.F. Wilson, N. Dunstan, et al. 2011. A combined long-range phasing and long haplotype imputation method to impute phase for SNP genotypes. Genet. Sel. Evol. 43(1): 12. doi: 10.1186/1297-9686-43-12.

Hickey, J.M., and A. Kranis. 2013. Extending long-range phasing and haplotype library imputation methods to impute genotypes on sex chromosomes. Genet. Sel. Evol. 45(1): 10. doi: 10.1186/1297-9686-45-10.

Howie, B.N., P. Donnelly, and J. Marchini. 2009. A flexible and accurate genotype imputation method for the next generation of genome-wide association studies. PLoS Genet. 5(6): e1000529.

Jacobson, A., L. Lian, S. Zhong, and R. Bernardo. 2014. General Combining Ability Model for Genomewide Selection in a Biparental Cross. Crop Sci. 54(3): 895. doi: 10.2135/cropsci2013.11.0774.

Jacobson, A., L. Lian, S. Zhong, and R. Bernardo. 2015. Marker imputation before genomewide selection in biparental maize populations. Plant Genome 8(2): 9. doi: doi:10.3835/plantgenome2014.10.0078.

Kong, A., G. Masson, M.L. Frigge, A. Gylfason, P. Zusmanovich, et al. 2008. Detection of sharing by descent, long-range phasing and haplotype imputation. Nat. Genet. 40(9): 1068–1075. doi: 10.1038/ng.216.

Li, Y., C.J. Willer, J. Ding, P. Scheet, and G.R. Abecasis. 2010. MaCH: using sequence and genotype data to estimate haplotypes and unobserved genotypes. Genet. Epidemiol. 34(8): 816–834. doi: 10.1002/gepi.20533.

Loh, P.-R., P. Danecek, P.F. Palamara, C. Fuchsberger, Y. A Reshef, et al. 2016. Reference-based phasing using the Haplotype Reference Consortium panel. Nat. Genet. 48(11): 1443–1448. doi: 10.1038/ng.3679.

O’Connell, J., K. Sharp, N. Shrine, L. Wain, I. Hall, et al. 2016. Haplotype estimation for biobank-scale data sets. Nat. Genet. advance online publication. doi: 10.1038/ng.3583.

Sargolzaei, M., J.P. Chesnais, and F.S. Schenkel. 2011. FImpute - An efficient imputation algorithm for dairy cattle populations. J. Dairy Sci. 94 (E-Suppl. 1): 421.

VanRaden, P.M., C. Sun, and J.R. O’Connell. 2015. Fast imputation using medium or low-coverage sequence data. BMC Genet. 16(1): 82. doi: 10.1186/s12863-015-0243-7.

